# Autocatalytic growth offset by constant degradation explains mass accumulation during B cell activation

**DOI:** 10.64898/2026.05.22.727310

**Authors:** Sophia M. Remick, Bipasha Mazumder, Koushik Roy, Thomas A. Zangle

## Abstract

How cells regulate growth is a fundamental question in cell biology. Here we use quantitative phase imaging (QPI) to measure the dry mass of individual naïve primary mouse B cells during *ex vivo* stimulation. B cells undergo an approximately 4-fold increase in mass prior to first division, far exceeding the doubling in size typical of immortalized cell lines and enabling discrimination of growth models that are otherwise indistinguishable. We find that a pure exponential model of mass accumulation exhibits systematic error that is resolved by an offset exponential model, indicating that only an active fraction of cell mass participates in biosynthesis. Monte Carlo simulations establish when these two models can be discriminated as a function of final to initial mass ratio, the magnitude of the degradative pressure, and measurement precision. Analysis of growth rate versus mass independently confirms the offset exponential model and implies that it can be interpreted as indicating a constant degradative pressure that opposes exponential biosynthesis. This model is structurally equivalent to super-exponential growth models derived from autocatalytic ribosome dynamics in bacteria, with the exponential rate reflecting autocatalytic accumulation of biosynthetic machinery and the active mass fraction serving as a proxy for initial ribosomal capacity. Here, the low active fraction in quiescent B cells (∼20%) is consistent with a limited ribosomal content of resting lymphocytes. Comparison of follicular and marginal zone B cells stimulated with CpG reveals a conserved degradative pressure across subtypes and stimulation conditions. These results demonstrate that deviations from pure exponential growth, previously identified in bacteria, extend to mammalian cells, suggesting that autocatalytic ribosome accumulation offset by constitutive protein turnover is a conserved feature of single-cell growth dynamics across domains of life.

## Introduction

Cell size regulation is fundamental to multiple biological processes, including tissue homeostasis, development, and the immune response (1). In immunity in particular, the size of a founder B cell at division can predict the maximum number of division cycles of the founder cell (2), influencing the strength of the immune response. However, the exact mechanisms underlying cell size regulation remain poorly understood. Measurements of the overall size of cells, and their rate of change, have provided key clues to these mechanisms. A central question is whether cells grow linearly, with a constant rate of mass addition (d*m*/d*t* = constant), or exponentially, with a mass-dependent rate (d*m*/d*t* proportional to *m*) (3).

Yeast and bacterial systems have provided key insights into growth regulation, due in part to the relatively simple geometry of typical unicellular organisms, which makes it easier to quantify cell size and growth. In fission yeast, a bilinear growth model has been proposed to fit cell length over time (4), while studies in budding yeast and bacteria have shown that growth rate is linked to ribosome content (5). Exponential growth is expected under conditions where ribosome availability is limiting, as the rate of biomass accumulation is then proportional to existing biomass (5).

Cell growth is a combined effect of the synthesis and degradation of cellular components. Recent work modeling growth of yeast and bacteria indicates that protein turnover plays a key role in regulating cell growth, and that this effect can be captured by modeling the fraction of ribosomes within cells as inactive, predicting a degradative pressure that limits cell growth (6). This model predicts deviation from a pure exponential model of cell growth (6), an effect that has been further explored in models coupling ribosome and amino acid pools (7). Furthermore, single-cell measurements of rod-shaped bacteria have revealed that growth within the cell cycle is super-exponential, with the instantaneous growth rate increasing over time, an effect that can be explained by autocatalytic ribosome production coupled to nonuniform cell elongation (8). In proliferating mammalian cell lines, sub-exponential mass-dependent growth rate modulation has been identified as a possible mechanism for maintaining cell mass homeostasis (9). Together, these studies suggest that deviations from pure exponential growth are a general feature of single-cell growth dynamics, with functional consequences for size homeostasis across organisms.

In mammalian systems, several approaches have been used to study growth regulation. Volume measurements during confined growth of immortalized cell lines suggest adder-like behavior, where a fixed volume is added per cell cycle independent of birth size (10). A coulter counter device that exploits the spherical geometry of detached cells has found evidence for size-dependent limits on cell growth (11). However, in addition to size, mammalian cells vary in density (12), potentially interfering with interpretation of results based on cell volume measurement alone.

Measurement of cell mass is potentially a more direct readout of biosynthetic output, as it reflects the sum of macromolecular accumulation through the cell cycle. A number of approaches are available to measure the mass of individual cells that are minimally affected by cell shape. For example, fluorescent labels that nonspecifically bind proteins can be used to assess population-average cell growth from fixed cells using flow cytometry (13). Buoyant mass measurements using suspended microchannel resonators offer ultra-high precision, with reported noise floors below 0.1% for single cells (14,15), but are limited to one cell per resonator, though multiple devices can be used in parallel (16).

Another approach to precisely measure cell mass is quantitative phase imaging (QPI), which measures the optical path length shift of light as it passes through cells (17,18). For a given cellular component, this phase shift is linearly proportional to mass, characterized by the specific refractive increment (19,20). When applied to whole cells, an appropriate weighted average specific refractive increment is assumed (21), enabling QPI measurements to be interpreted as measurements of whole-cell dry mass (22). QPI has been applied to monitor changes in growth through the cell cycle (23), during cell death (24), and in response to specific growth inhibitors (25). The typical precision of QPI, after accounting for errors in the raw phase measurement as well as image processing operations required to measure cell mass, is approximately 1% in cell mass (26,27). Though this is not as precise as the typical performance of microchannel resonators, QPI enables longitudinal measurements of thousands of single cells simultaneously (25), enabling high-throughput studies of cell growth. We have previously applied QPI for lineage tracing in naïve, mouse, primary B cells, revealing that select cells show a dramatic, > 5-fold increase in mass with apparent super-exponential growth prior to cell division (28), making naïve B cells an ideal system to study growth regulation.

Here, we apply QPI for measurement of naïve primary B cells during *ex vivo* stimulation, leveraging the fact that B cells grow substantially upon stimulation. These data indicate that a pure exponential model has systematic error that can be explained by an additional constant term in fits of cell mass over time. Statistical modeling reveals the interplay between measurement precision and the ability of a growth assay to distinguish this offset exponential model from pure exponential growth, indicating a possible explanation for why this feature has not been reported previously in mammalian cells. We show that this constant term corresponds to a degradative pressure on cell growth that acts throughout the cell cycle, analogous to that which has been observed in bacteria and yeast (6,8). This degradative pressure does not vary significantly with stimulation condition or B cell subtype, providing evidence of a near-constant degradative drag on cell mass accumulation in primary B cells.

## Results

Primary mouse B cells were stimulated with CpG oligonucleotides to induce cell growth and expansion and typically divide in ∼1-2 days post stimulation. As shown in **Fig. 1a** this division is preceded by a large expansion in overall cell size. This individual cell increases in dry mass from 40 pg at the start of the experiment, to 174 pg just prior to cell division, a 4.4-fold increase in mass (**Fig. 1b**). This is typical of this population of B cells which displayed a of 4.6+/- 1.6-fold increase in mass prior to the first cell division (average +/- standard deviation, *n* = 1899, **Fig. S1**), and significantly larger than the doubling (2-fold) increase in mass typically observed in immortalized cell lines through the cell cycle. With this large increase in mass, combined with the high precision of QPI, the exponential curvature of *m*(*t*) is clearly observed. However, fitting with a pure exponential model results in an underestimate at the start and end of the observation period (**Fig. 1b**, dark yellow line, R^2^ = 0.95). Modifying the model to include a constant term dramatically improves the fit (**Fig. 1b**, blue line, R^2^ = 0.99). Akaike information criteria (AIC) scores balance decreases in goodness of fit against an increase in number of parameters. Comparison of the two plots using the AIC indicates that the offset exponential model is strongly preferred (*p* < 1e-40). A plot of the residual to the fit versus time clearly shows that the addition of a constant term improves the exponential fit to *m*(*t*), particularly at the start and end of the measurement period (**Fig. 1c**). The average ratio of *C*/(*A*+*C*) from least squares nonlinear fitting is 0.79 +/- 0.04, indicating that the constant, *C*, is a significant fraction of cell initial mass, *A*+*C*, though this has little impact on the total increase in mass, *m*_*f*_/*m*_*0*_ (**Fig. 1d**).

**Figure 1.**
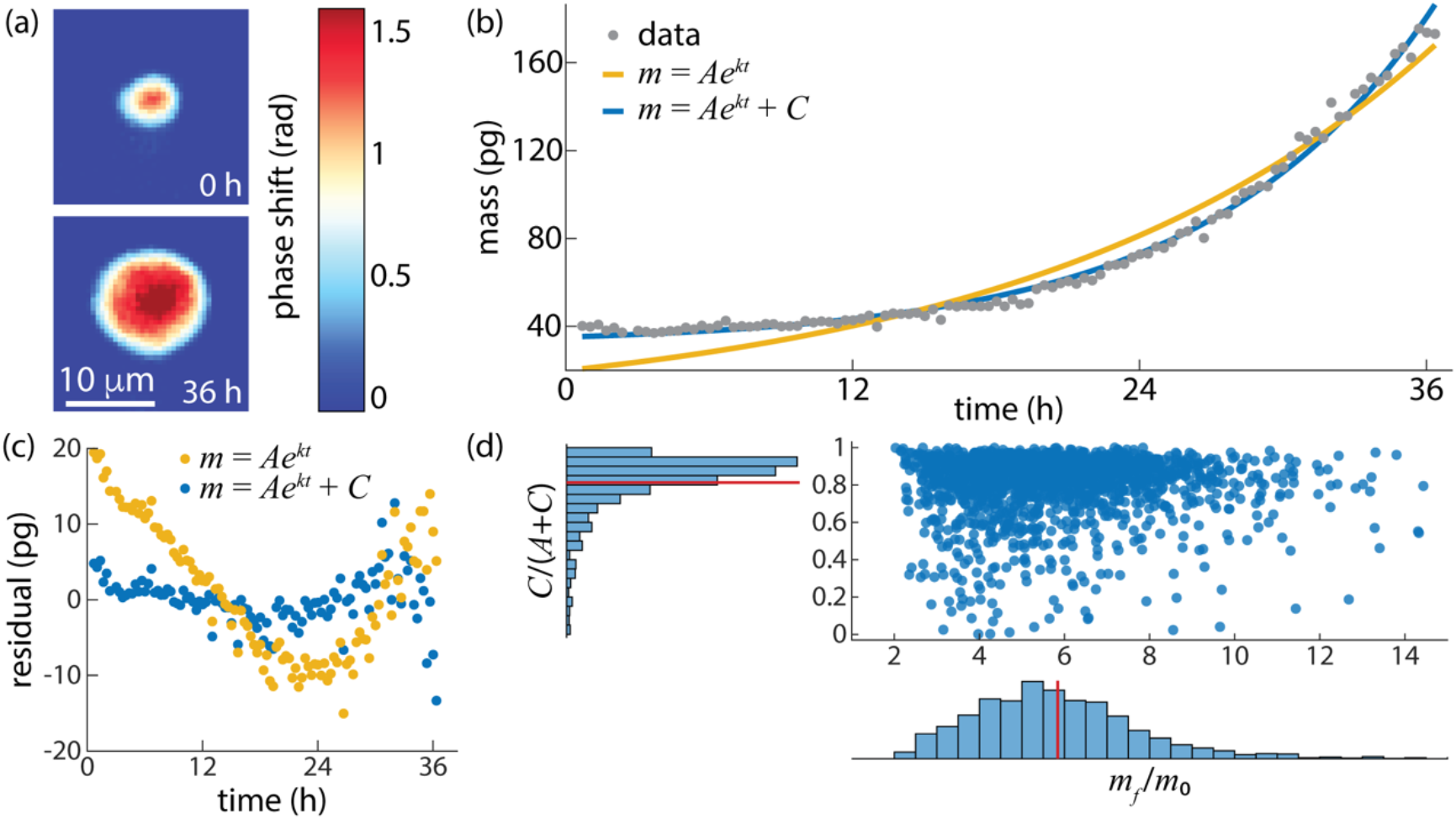
Mass over time of primary B cells fits an offset exponential model. (a) QPI image of naïve primary mouse marginal zone B cell at *t* = 0 h and *t* = 36 h post CpG stimulation showing an increase in cell size. Colormap indicates mass, dark blue is background, red shows regions of high mass. (b) mass over time (grey) for the cell shown in (a). Best fit lines (least squared error) for a pure exponential (yellow) and offset exponential (blue) are shown. (c) Residuals to the pure exponential and offset exponential models for data in (b). (d) Distributions of individual B cell final (mass at division) over initial mass (*m*_*f*_/*m*_*0*_, *x*-axis) and ratio of model constant (*C*) to initial mass from model fitting (*A*+*C*). Mean values shown as solid red lines on histograms. *n* = 1899 cells.

In measurements of immortalized cells that divide after approximately doubling in mass, the addition of a constant to the pure exponential mass accumulation model is not typically justified from fitting mass as a function of time. To test when the exponential and offset exponential models can be discriminated, we performed a series of Monte Carlo simulations to fit artificial cell data across the experimentally observable range of model parameters (**Fig. 2a**). These simulation data indicate that the difference between the exponential and offset exponential models depends strongly on the ratio of final to initial mass (**Fig. 2b**). For the typical doubling of cell mass during the cell cycle the two models are nearly indistinguishable by SSE, and visually very similar (**Fig. S2**) with a small amount of systematic variation over time. However, at a mass ratio of greater than 3, as observed frequently (89%) in primary B cells (**Fig 1d**), the difference between the two models is pronounced.

**Figure 2.**
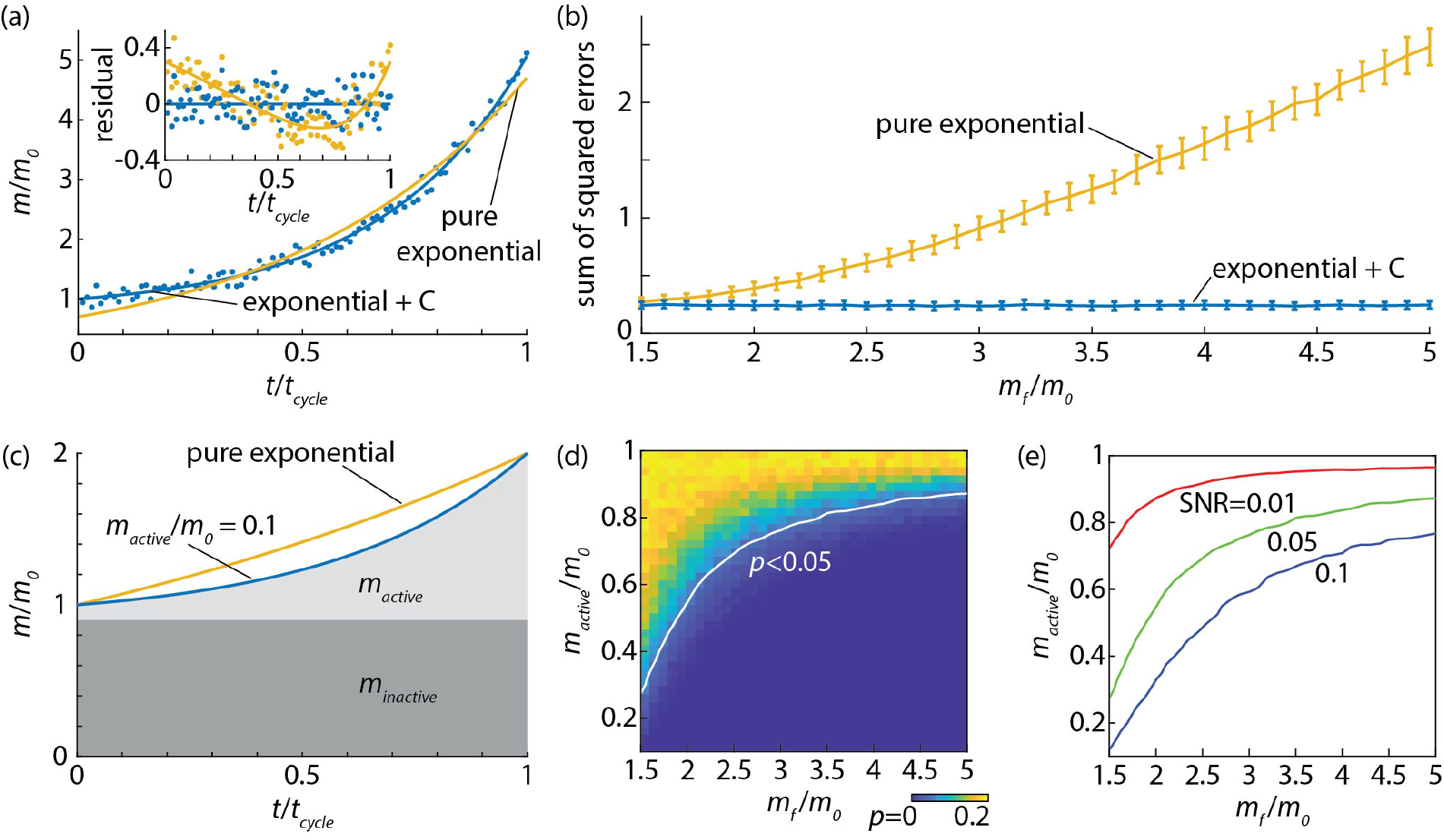
Impact of model parameters on fit of offset exponential model. (a) Simulated normalized mass versus time data for a cell following an offset exponential growth trend to 5-fold initial mass with *m*_*active/*_*m*_*0*_ = 0.2 and a measurement signal to noise ratio (SNR) of 0.1. Data shown in blue with best fit offset exponential as a solid blue line, and best fit pure exponential in yellow. Inset shows the residual for both fits (points) and the difference between the fit and simulated data without noise as solid lines. (b) Sum of squared errors for curve fitting (*m*_*active/*_*m*_*0*_ = 0.2, SNR = 0.05) versus ratio of final to initial mass. Error bars show standard deviation from 100 simulated cells per condition. (c) Comparison of pure exponential and offset exponential (*m*_*active/*_*m*_*0*_ = 0.1) growth curves. The shaded areas indicate the ‘inactive’ and ‘active’ components of cell mass in the offset exponential model. (d) Probability, *p*, that the simpler (exponential) model fits data better than the more complicated (offset exponential) model from Akaike information criteria (AIC) for SNR = 0.05 from results for 100 simulated cells at each condition. The dark blue region in the lower right (high *m*_*f/*_*m*_*0*_, low *m*_*active/*_*m*_*0*_) indicates a significantly better fit for the offset exponential model. (e) Iso-curves of *p* = 0.05 from the AIC test or three values of SNR, 0.01, 0.05, and 0.1.

Another key parameter in the model is the portion of a cell’s initial mass that contributes to exponential growth. This appears in normalized form as *A*/(*A+C*), with *A*/(*A+C*) = 1 corresponding to a pure exponential. A decrease in *A*/(*A+C*) results in a deeper curvature to plots of mass versus time larger and a larger deviation from a pure exponential (**Fig. 2c**). Consistent with this, primary B cell data show a preferred fit to the offset exponential model at lower *A*/(*A+C*) values (**Fig. S3**). This is exaggerated by larger ratio of final to initial mass (**Fig. S4a**). This parameter, *A*/(*A+C*) can alternately be interpreted as the fraction of cell mass that contributes to exponential growth, e.g. *m*_*active/*_*m*_*0*_, so that, for example, to achieve an equivalent mass increase over the cell cycle period, the corresponding value of the exponential constant, *k*, increases (**Fig. S4b**).

The ability to distinguish the exponential and offset exponential models is, therefore, a function of both the increase in cell mass over the cell cycle or observation period, *m*_*f*_/*m*_*0*_, as well as *A*/(*A+C*). To capture this, we used Monte Carlo simulations of artificial cell data over a range of these two parameters and evaluated goodness of fit. As expected, the AIC probability shows an increasing difference between the two models as *m*_*f/*_*m*_*0*_ increases and *A*/(*A+C*) decreases (**Fig 2d**). The location of the boundary between the two models is significantly affected by the underlying measurement SNR (**Fig 2e**), with lower SNR increasing the region in which the offset exponential model is 95% preferred, even for *m*_*f*_/*m*_*0*_ = 2. For SNR = 0.01, corresponding to a 1% error in mass which is about the lower limit reported for cell measurements with QPI (26,29), the difference between these models should be distinguishable at *m*_*active/*_*m*_*0*_ values as high as 0.8 (or inactive mass of only about 20% of initial cell mass). We observe similar results with an F-test that compares these two nested models by estimating whether the extra parameter in the offset exponential model reduces the SSE by more than expected from random chance (**Fig S5**).

The addition of a constant to the pure exponential model for mass accumulation also appears in measurements of the instantaneous rate of mass change, *dm*/*dt*, as a function of *m*. Starting with the observed form of mass as a function of time:

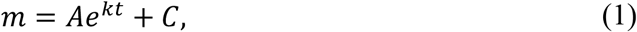

yields the following for the derivative of mass over time, *dm*/*dt*:

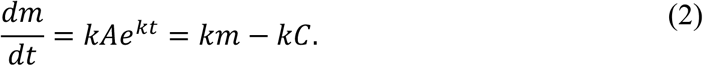

Measurement of the rate of change has previously been used to show the difference between linear and exponential models (3). The offset exponential model predicts that *dm*/*dt* will be linearly related to *m* with a negative intercept (Equation 2). In contrast, pure exponential growth implies that *dm*/*dt* is linearly proportional to *m* with an intercept of 0. Therefore, for a positive value of *C*, as observed in the cell shown in **Figure 1a-c**, we anticipate a non-zero intercept equal to *-kC* when plotting the instantaneous rate of mass change versus mass, as we observe for this cell (**Fig 3a**). Here, the value of *C* estimated from a best fit to *dm*/*dt* versus *m* (**Fig 3a**) is 32.0 +/- 1.8 pg (95% confidence interval), which agrees well with the value of 31.9 +/- 2.8 pg (95% confidence interval) determined from directly fitting *m*(*t*) (**Fig 1b**). Across *n* = 1899 founder B cells, these two methods of determining *C* are generally in agreement (**Fig 3b**, R^2^ = 0.90). Additionally, linear regression to all instantaneous *dm*/*dt* estimated from individual B cells stimulated with CpG has a negative intercept (**Fig 3c**), predicting a population *C*_*dm/dt*_ = 18.6 +/- 0.2 pg, slightly less than the overall average from fitting *m*(*t*) of *C*_*m*_ = 22.2 +/- 0.6 pg, possibly due to the delay of imaging to 40 minutes post stimulation, causing a slight increase in mass at *t*_*0*_ from *m*(*t*) data.

**Figure 3.**
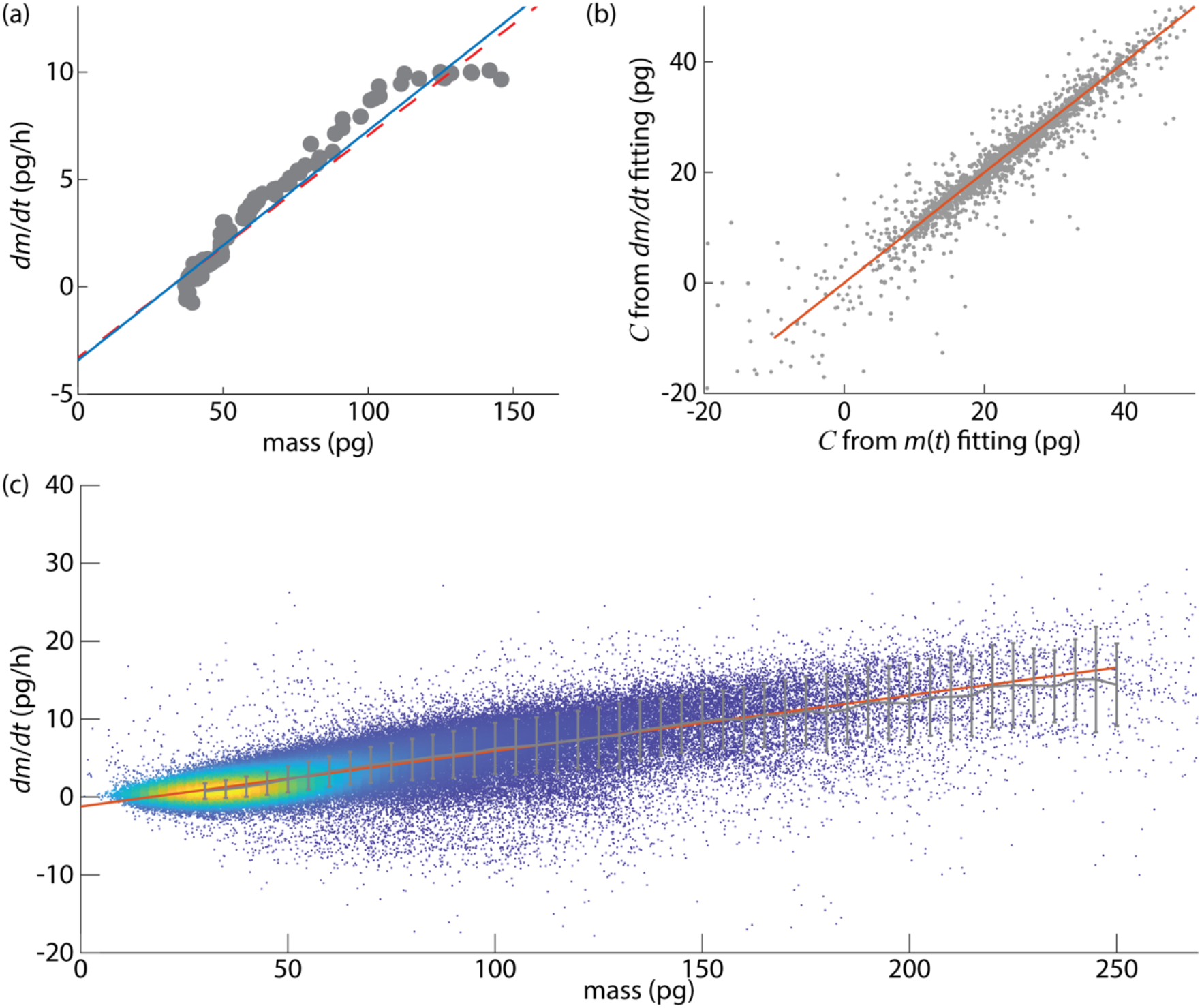
Offset exponential model predicts a negative degradative rate at the limit of zero cell mass. (a) Growth rate, measured as instantaneous *dm*/*dt*, versus mass for the cell data shown in **Figure 1**. Least squared best fit line shown in solid blue. Estimated relationship from fitting to mass over time data shown as a dotted red line. (b) Comparison of *C* in offset exponential model estimated from linear fitting of instantaneous *dm*/*dt* vs. mass compared to C determined from fitting the offset exponential model to *m*(*t*) data. A line of *C*_*dm/dt*_ = *C*_*m*_ (the theoretical value) is shown in red. (c) Instantaneous *dm*/*dt* vs. mass from all CpG stimulated wild type B cells. b,c: *n* = 1899 cells.

We observe a moderate correlation (0.6) between initial and final mass (**Fig 4a**) and a low correlation (0.06) between the exponential growth rate, *k*, and final mass (**Fig 4b**), suggesting that ultimate cell size in naïve B cells, which is linked to proliferation capacity (2), is mostly related to initial size, not cell-specific growth rates. However, the inverse of growth rate, 1/*k*, is correlated with active mass fraction, *A*, (**Fig 4c**, R = 0.73), suggesting that larger growth rate acts to compensate for reduced initial biosynthetic capacity, rather than directly determining cell size.

**Figure 4.**
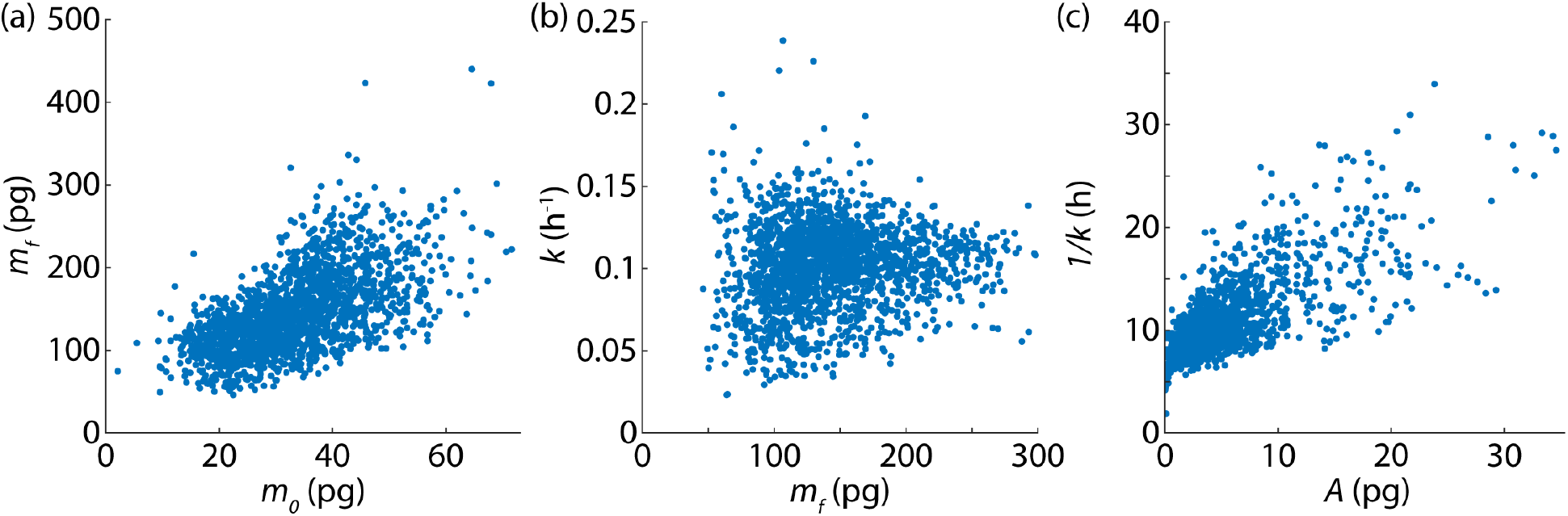
Growth rate of naïve B cells is not related to ultimate size and compensates for reduced initial biosynthetic capacity. (a) Observed final mass, *m*_*f*_ versus initial mass, *m*_*0*_, of CpG stimulated naïve B cells. (b) Exponential growth constant, *k*, (e.g. Equation 1) from fitting the offset exponential model versus final mass, *m*_*f*_. (c) Inverse of exponential growth constant versus initial active mass, *A*, from fitting the offset exponential model. *n* = 1899 cells.

To investigate the relationship between the observed exponential growth constant and the degradation rate predicted by the offset exponential model, we can look at the rate of degradation, computed as *kC* (last term in Equation 2). Normalizing by initial mass, *A*+*C*, yields a degradative rate for the cell at the start of the cell cycle:

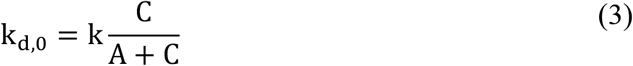

Where *k*_*d,0*_ is the degrative rate constant at *t* = 0. Plotting *k*_*d,0*_ versus *k* yields a positive correlation as expected for a positive constant, *C* (*R* = 0.97, **Fig. 5a**). The *x*-intercept of this relationship is 0.029 +/- 0.001 h^-1^, indicating a limit of positive growth rate at zero degradation rate. We can then define an overall growth rate, *λ*, consistent with previous modeling of degradative rates in *E. Coli* and *S. cerevisiae* ADDIN EN.CITE (6), in terms of the offset exponential model:

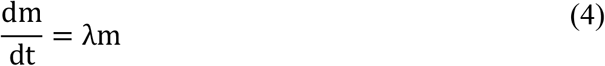

**Figure 5.**
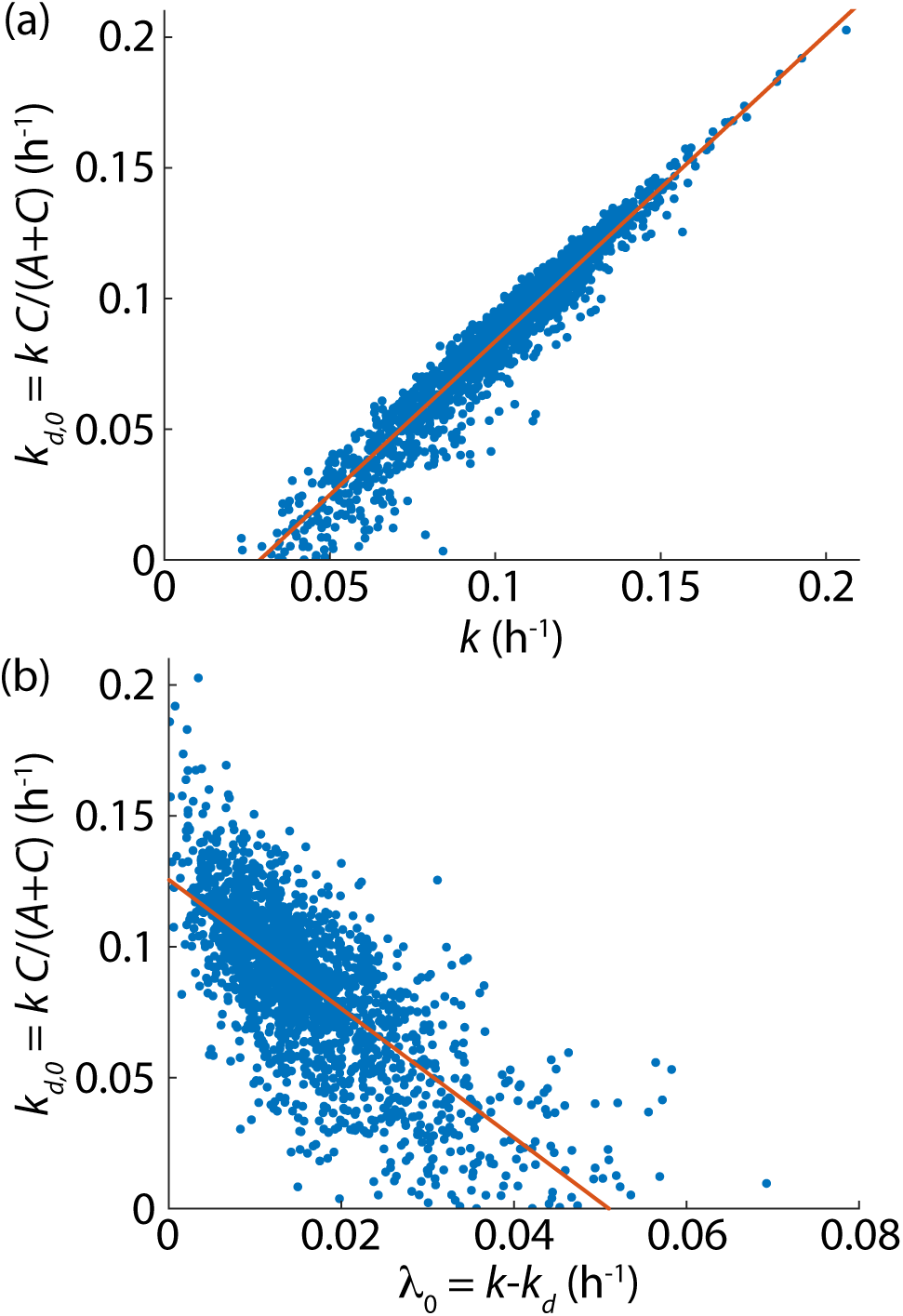
Offset exponential model predicts limiting degradative rate in primary B cells is comparable to overall growth rates. (a) Degradative rate at *t* = 0, *k*_*d,0*_, versus exponential rate constant, *k*. Linear least squares best fit line shown in red. (b) Degradative rate at *t* = 0 versus apparent exponential growth rate at *t* = =, *λ*_*0*_. Linear least squares best fit line shown in red. *n* = 1899 cells.

where *λ* is the apparent exponential growth rate of the cell, or overall growth rate normalized by mass at a given time. Note that, for a cell growing as predicted by the offset exponential model (Equation 1), this implies *λ* must be a function of time. Combining equations 2 and 4 yields:

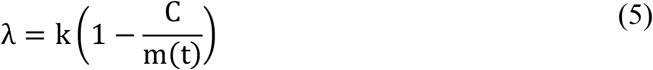

Normalizing by the mass at time 0, which is equal to A+C in the model (Equation 1) yields the following expression for the initial apparent exponential growth rate, *λ*_*0*_:

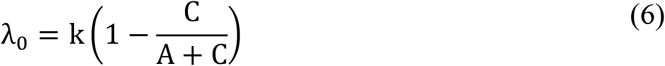

We find that *λ*_*0*_ decreases with increasing *k*_*d,0*_ (**Fig 5b**) as observed previously in *E. Coli* and *S. cerevisiae* (6). The estimated degradation rate at zero growth, or value of *k*_*d,0*_ at *λ*_*0*_ = 0, is 0.126 +/-0.002 h^-1^, which is comparable to the overall observed exponential growth constant (**Fig 4**), supporting the idea that degradation plays a large role in determining the growth of naïve B cells. These findings are consistent when the data is analyzed for B cells separately as well. The offset exponential model was preferred for both subtypes and the initial degradation rate, *k*_*d,0*_, did not differ significantly across conditions (**Fig. S6)**.

## Discussion

The offset exponential model of cell mass accumulation has not been previously proposed for mammalian cells, likely because the model is difficult to distinguish from pure exponential growth at the typical doubling of mass observed in immortalized cell lines (**Fig. 2**). Distinguishing even between linear and exponential growth models with typical data can be challenging (3). Two features of our experimental system enable this distinction. First, the approximately 4-fold increase in mass of primary B cells upon stimulation exaggerates the difference between the exponential and offset exponential models (**Fig. 1**). Second, single-cell mass measurements are essential, as any monotonic mass accumulation pattern in populations or multi-cellular clusters will tend towards a pure exponential as the number of cells increases (30).

The offset exponential model presented here has two complementary interpretations. In terms of mass as a function of time (Equation 1), the model implies that a portion of cell mass does not contribute to the generation of additional biomass. We term this the “inactive” mass fraction. This fraction can be substantial, comprising approximately 80% of initial mass in naïve primary B cells. In this interpretation, the “active” portion of cell mass performs proportionally more biosynthetic work than in a corresponding pure exponential model, as reflected by the increase in the exponential rate constant *k* relative to the pure exponential model (Fig. S4). The alternative interpretation, suggested by the analysis of d*m*/d*t* versus mass (Fig. 3), is that the constant term represents a degradative pressure that opposes biosynthesis at a fixed rate (Equation 2). This interpretation is supported by recent theoretical work in yeast and bacteria demonstrating the importance of incorporating protein degradation in models of cell growth (6).

Recent modeling of ribosomal dynamics points towards a resolution of these two views. In bacteria, the observation that single-cell growth within the cell cycle is super-exponential has been explained by a model in which biomass accumulation is proportional to ribosome abundance, and ribosomes are autocatalytic (8). This yields a growth equation for bacterial cell length, *L*, of the form d*L*/d*t* = *k*(*L* – λ), where λ represents a non-growing portion of the cell. This is structurally equivalent to the offset exponential model (Equation 2), with the constant *C* = *k*λ representing the mass that does not participate in ribosome-driven growth. In this framework, the exponential rate *k* extracted from the growth rate analysis (**Fig. 3**) directly reflects the rate of autocatalytic accumulation of biosynthetic machinery, while the active mass fraction *A*/(*A*+*C*) provides an experimentally accessible proxy for the initial ribosomal capacity of a cell relative to its total mass. In quiescent B cells, this fraction is low (approximately 0.2 on average), indicating a low ribosomal content expected of resting lymphocytes. As ribosomes accumulate autocatalytically following activation, the fraction of mass engaged in biosynthesis increases, producing an accelerating apparent growth rate (Equation 5). Additionally, recent measurements of yeast growth, mRNA concentration, and rRNA sequencing combined with growth modeling suggests that both inactive mass fraction as well as degradation may play a term in growth of eukaryotic cells (31). The observation that bacterial, yeast, and mammalian cells deviate from pure exponential growth in a manner consistent with autocatalytic ribosome dynamics suggests that this may be a conserved feature of cellular growth across domains of life, with the magnitude of the deviation reflecting the initial mismatch between biosynthetic capacity and total cell mass.

While single cell mass versus time data provide the clearest evidence for the growth model shown here (**Fig. 1**), measurement of instantaneous growth rate versus mass (**Fig. 3**) may prove useful for extending these findings to other cell types that do not exhibit the large mass increase seen during B cell activation. Because this approach relies on the slope and intercept of a linear fit (Equation 2) rather than distinguishing the curvature of mass as a function of time, it is potentially more sensitive to deviations from pure exponential growth. However, this approach does not directly reveal the initial active mass fraction which requires fitting the full mass over time trajectory.

The balance between synthesis and degradation during B cell activation has important biological implications. Excessive anabolism has been shown to lead to metabolic stress, excess B cell growth, and enhanced sensitivity to apoptosis following pre-B cell receptor stimulation (32). Unrestrained synthesis is detrimental to B cell development at the large pre-B cell stage, which serves as a checkpoint for self-antigen regulation to prevent autoimmunity (33), and excess synthesis of cellular components without degradation could contribute to the generation of autoimmune responses (34). Furthermore, the size of a founder B cell at the time of first division can predict the number of subsequent division cycles (2), suggesting that excessive synthesis without compensatory degradation during B cell activation could lead to excessive proliferation. Consistent with this, diffuse large B cell lymphoma, one of the fastest growing B cell lymphomas (35), features cells at least twice the size of normal B cells (36,37). Our observation that higher growth rates correlate with higher degradation rates (**Fig. 5**) suggests that degradation may serve as a homeostatic mechanism to restrain growth during B cell activation, with implications for understanding immune responses.

Overall, this work provides evidence that protein degradation plays a significant and quantifiable role in the growth of naïve primary B cells, and that this degradative pressure can be captured by a simple constant term appended to the standard exponential growth model. The quantitative framework developed here, combining single-cell QPI measurements with Monte Carlo analysis of model discrimination, provides a roadmap for identifying similar deviations from pure exponential growth in other mammalian cell types and points towards methods to quantify degradative rates in mammalian cells more generally.

## Methods

### Quantitative phase imaging

Imaging was performed using a custom QPI microscope based on differential phase contrast (DPC) (38,39). The sample was imaged using a 10x objective with a numerical aperture of 0.25 placed 37 mm below a 27.6 mm square LED array. Images were captured by sequentially illuminating semicircular patterns in four directions the LED for DPC reconstruction. 12 locations were imaged within each well. Images were taken at 20 minute intervals for 50 hours, starting 40 min after stimulation to allow for temperature equilibration of the imaging plate and setting microscope focus positions.

### Image processing

Following phase reconstruction, raw intensity images underwent background correction, and cell segmentation was performed using CellposeSAM (40,41). After segmentation, cells were tracked by minimizing the displacement in x and y position as well as mass (42) to generate continuous mass vs. time tracks for each cell. To isolate founder cells, a filter was applied to only account for cells whose tracks were only present at frame 1 of the experiment. Data filtered on *m*_*f*_/*m*_*0*_ >2 and length > 20 frames. For measurement of instantaneous rate of mass change from mass over time data we computed the difference between mass measurements at sequential timepoints, divided by the time between frames, and smoothed with a Gaussian filter (σ = 5 frames).

### Primary B Cell Isolation

MZ (marginal zone) and FO (follicular cells) B cells were isolated from wild type C57BL/6 mouse spleens via negative selection with CD43 (Ly-48) MicroBeads (Miltneyi Biotec, 130-049-801), LS columns (Miltneyi Biotec, 130-042-401), and a MidiMACS separator (Miltneyi Biotec).

### Cell culture and dosing

B cells were cultured using RPMI (Gibco, 11875-119) plus 5 mM L-glutamine, 20 mM HEPES, β-ME (55 µM), 10% FBS, and penicillin–streptomycin. To promote cell adhesion, a flat bottom 96-well plate (Eppendorf, tissue culture treated) was coated with Poly-D-Lysine (Gibco). Each well received 50 μL of Poly-D-Lysine and the plate was incubated at room temperature for 1 hour. Following incubation, the coated wells were washed 3x with sterile DI water and left to dry for 2 hours. Cells were then seeded at a density of 5,000 cells/well in 200 μL culture media. Both MZ and FO cell types were treated with 250 nM CpG (Invivogen, tlrl-1668). Conditions were plated in triplicate wells.

### Monte Carlo curve fitting

Mass versus time data were fitted as a non-dimensionalized model, with cell mass, *m*, normalized by initial mass, *m*_*0*_, and time, *t*, normalized by the overall cell cycle time, *t*_*cycle*_. Individual *in silico* cell data was simulated by sampling from a normal distribution using randn() in Matlab (Mathworks). The signal to noise ratio (SNR) was defined as the ratio of the standard deviation of this distribution to the simulated initial cell mass. These simulated data were then fit by nonlinear least squares fitting in Matlab (lsqnonlin) and initial values of *m*_*active*_ = 1, *k* = 1, and *m*_*inactive*_ = 0 (for offset exponential model) over a parameter range of *m*_*active*_*/m*_*0*_ from 0.1 to 1, and *m*_*f*_*/m*_*0*_ from 1.5 to 5 and for three values of SNR, 0.01, 0.05, and 0.1 for a total of 400,000 simulated cells.

### Statistics

The difference in AIC values between the two model fits was computed as:

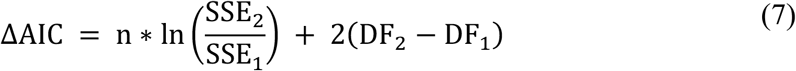

Where *n* is the number of measurement points, *DF* is the degrees of freedom for each model, model 1 is the simpler (pure exponential) model and model 2 is the more complex (offset exponential) model. The probability that the model 1 fit is preferred to model 2 is then:

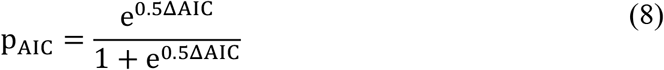

95% confidence intervals to fitted parameters were computed using cftool in Matlab (Mathworks).

The F-test statistic was computed using the fcdf function in Matlab (Mathworks).

## Notes

### Competing Interest Statement

The authors have declared no competing interest.

### Funding

- University of Utah

