## Supplementary material for "Autocatalytic growth offset by constant degradation explains mass accumulation during B cell activation": Supplemtary Information

**Contents**

Supplemental Figures S1-S6

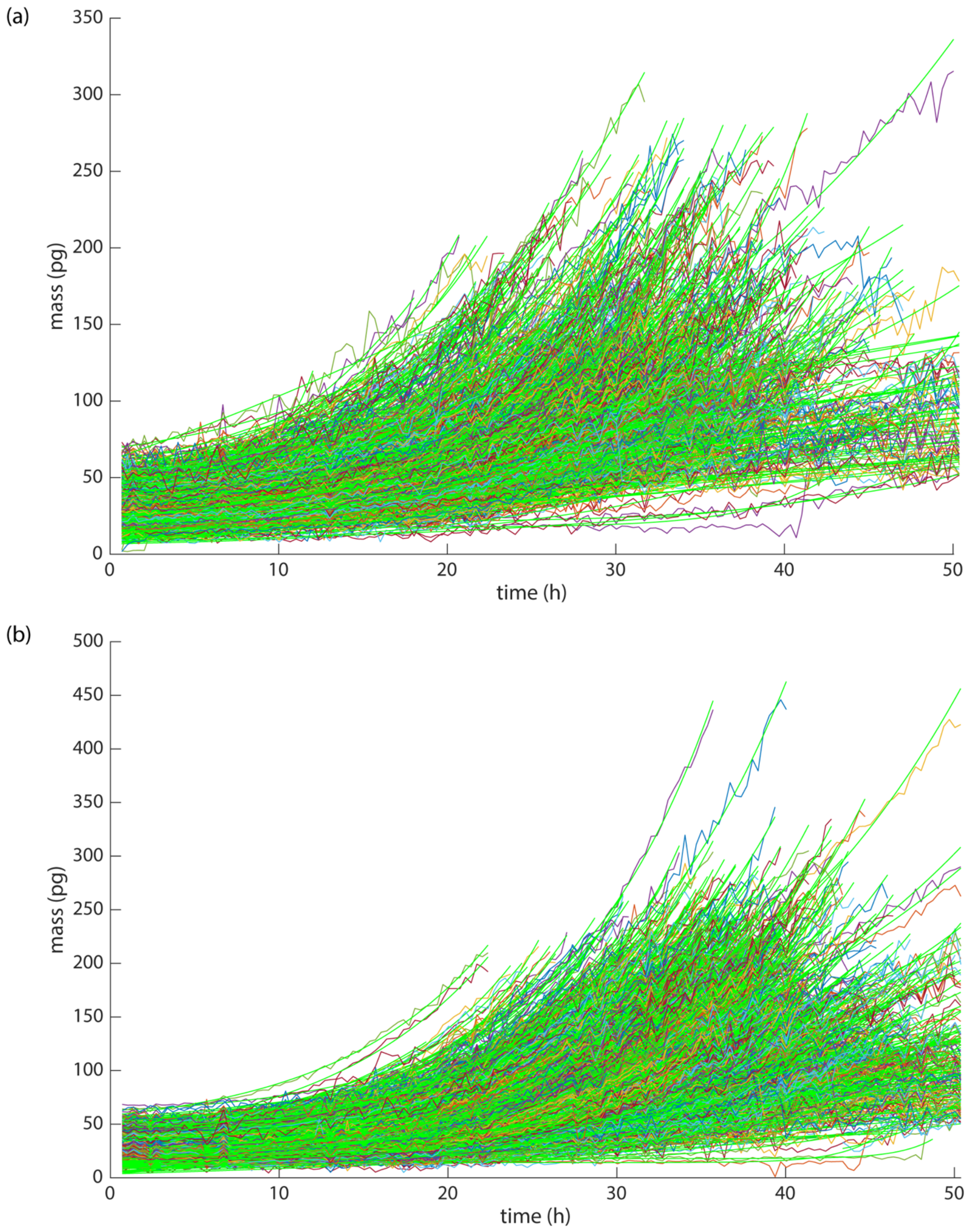


**Figure S1. Raw mass and offset exponential fits to wild type B cells stimulated with CpG.** Offset exponential fit lines shown in green. (a) $n$ = 509 marginal zone cells, (b) $n$ = 1390 follicular cells.

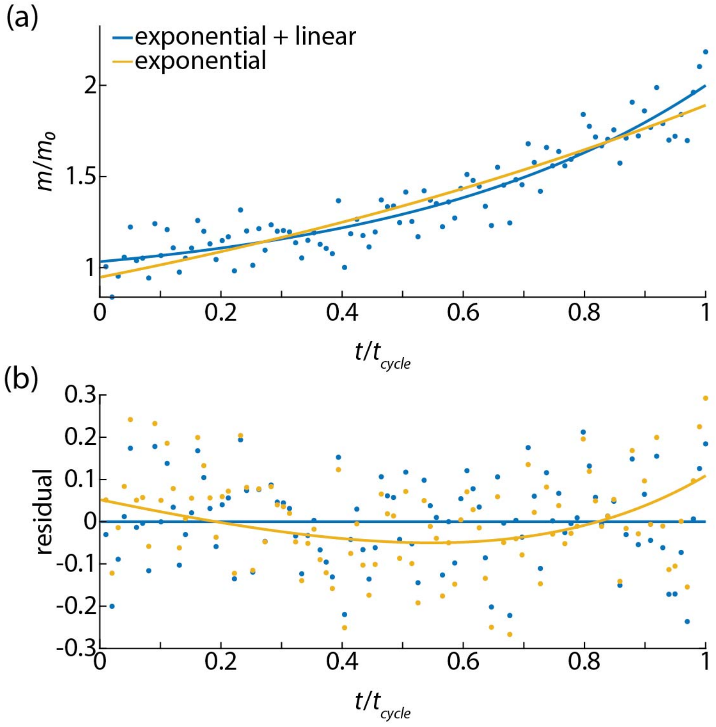


**Figure S2. Simulated data for $m_f/m_0 = 2$.** (a) Exponential and offset exponential fits to synthetic data with $m_f/m_0 = 2$, $m_{active}/m_0$ offset exponential as a solid blue line, and best fit pure exponential in yellow. (b) Residuals from fits in panel (a) shown as point and the difference between the fit and simulated data without noise shown as solid lines.

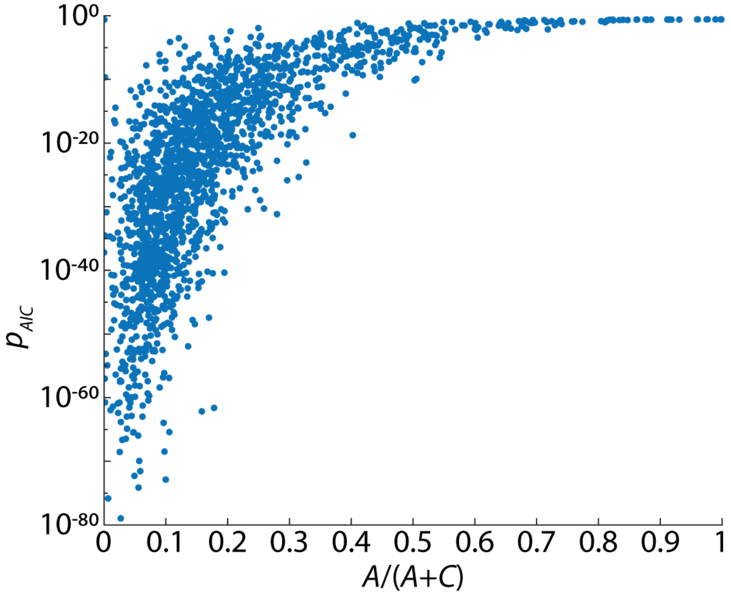


**Figure S3. Dependence of $p_{AIC}$ on $A/(A+C)$.** Probability the exponential model is preferred over the offset exponential model ($p_{AIC}$) versus ratio of A/(A+C) from $n$ = 1118 primary, wild-type B cells stimulated with CpG.

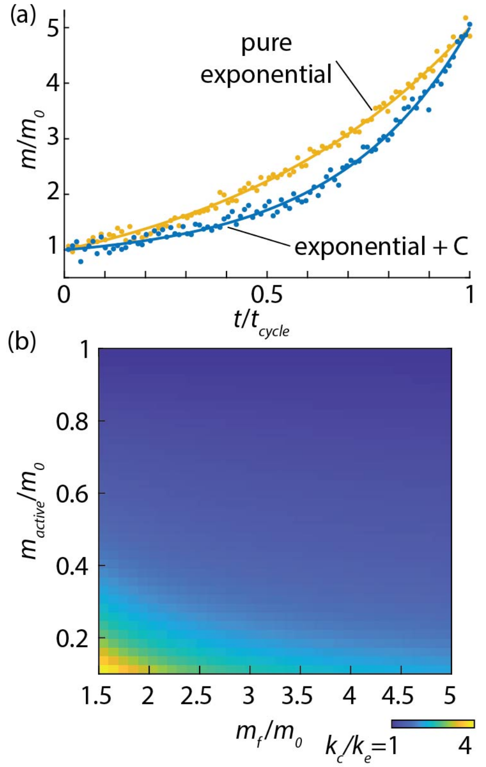


**Figure S4. Difference between model parameters in exponential and offset exponential models**. (a) Simulated mass versus time for exponential and offset exponential models at $m_f/m_0$ = 5 and $m_{active}/m_0$ = 0.2. simulated data points show SNR = 0.1. (b) Ratio of exponential coefficient, $k_c$, from offset exponential model relative to pure exponential model, $k_e$ showing increase as active mass fraction and overall increase in mass during the cell cycle decrease.

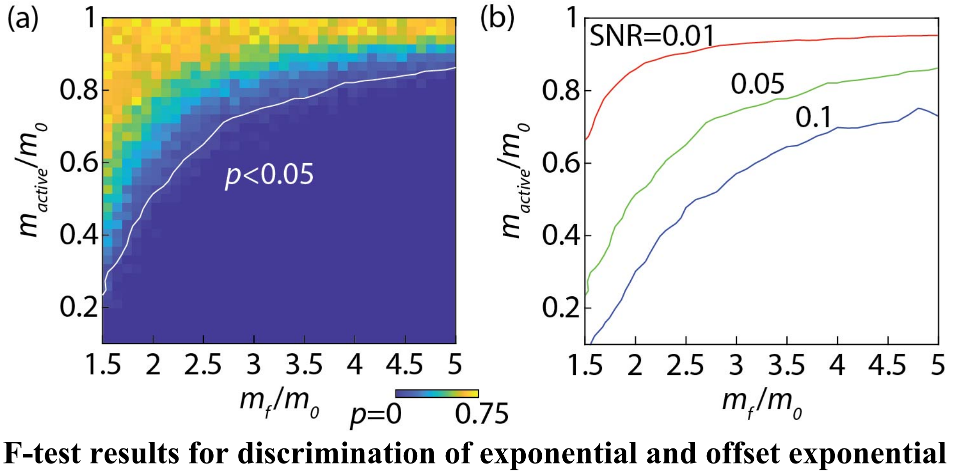


**Figure S5. F-test results for discrimination of exponential and offset exponential models.** (a) p-value from F-test results for simulated cell data. (b) Iso-curves of $p$ = 0.05 for three values of SNR, 0.01, 0.05, and 0.1.

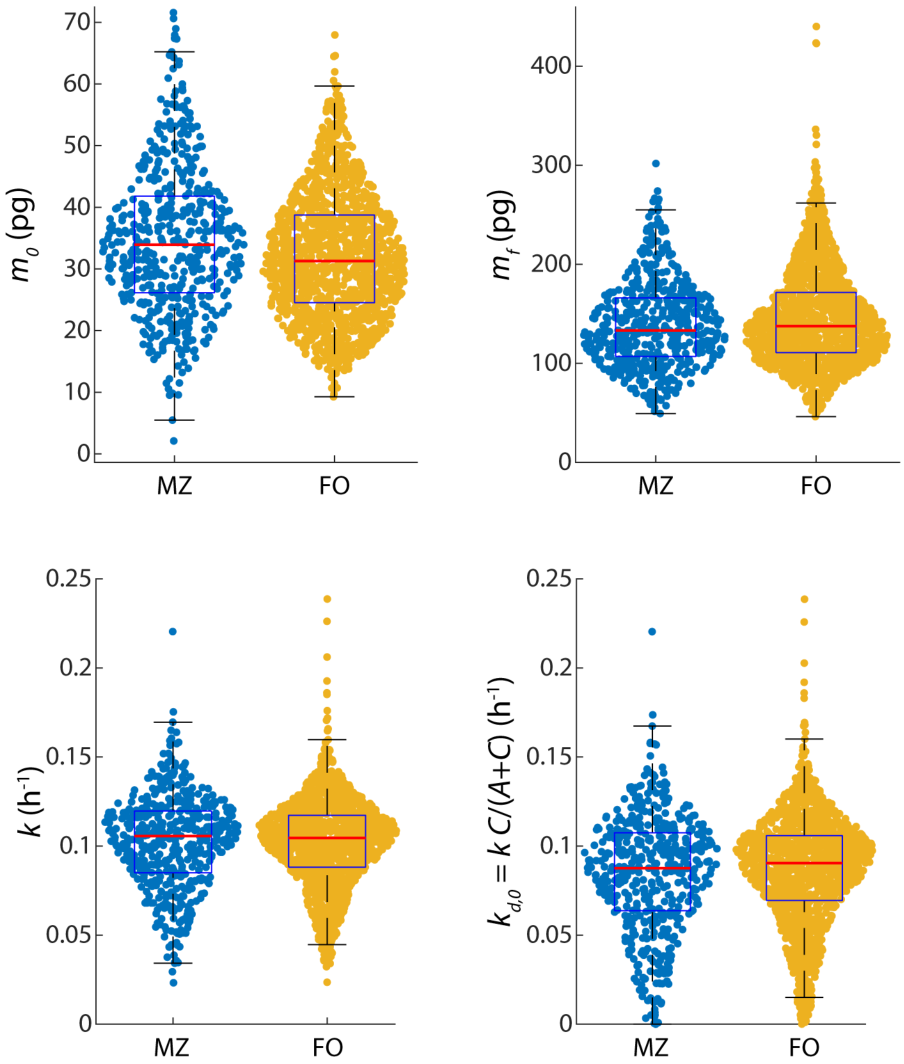


**Figure S6.** Comparison of offset exponential model parameters across B cell subtypes. Initial mass ($m_0$), final mass ($m_f$), exponential growth rate ($k$), and initial degradation rate ($k_{d,0}$) for follicular (FO) and marginal zone (MZ) B cells stimulated with CpG. Individual data points and boxplots (median shown in red) are shown for all conditions. $n = 509$ marginal zone cells and 1390 follicular cells.
